# On modeling ovarian aging and menopause timing

**DOI:** 10.1101/2025.04.07.647646

**Authors:** Sean D. Lawley, Nanette Santoro, Joshua Johnson

## Abstract

Mathematical modeling of ovarian aging and menopause timing has a long history, dating back a half-century to the models of Nobel Prize winner Robert G. Edwards. More recently, such models have been used to investigate clinical interventions for women, which underscores the importance of scientific rigor in model development and analysis. In this paper, we analyze a recent model published in the biophysics literature. We first correct an error which invalidates claims about menopause age in different populations. We then use stochastic analysis to show how this model is a reparameterization of a prior model and put it in the framework of several prior models, which enables the application of extreme value theory. We prove some general extreme value theory results and use them to obtain detailed estimates of menopause age in this model. In particular, we derive a new expected menopause age formula which is orders of magnitude more accurate than the previous heuristic estimate. We further obtain rigorous analytical estimates of the full menopause age distribution and all its moments. We conclude by using these mathematical results to elucidate the physiological sources of menopause age variability.

## 1 Introduction

The ovary is the organ responsible for producing gametes for reproduction in females. In many organisms, including mammals, the ovary also produces key hormones that participate in the control of reproductive cycles (in humans, menstrual cycles) and support the function of other organs throughout the body. Because ovarian follicles that contain egg cells (oocytes) are responsible for the production of these hormones, their presence is required for ongoing menstrual cycles.

Girls are born with around 10^5^ to 10^6^ dormant “primordial” ovarian follicles. Primordial follicle numbers decline during postnatal life, due to their commitment to growth and either survival to monthly ovulation, or, death. Loss of primordial follicles occurs over time, and eventually the number remaining is too low to give rise to numbers of growing follicles needed for ongoing ovarian function and reproductive cycling [1–3]. When menstrual cycles have been absent for a year, menopause is said to have been reached [3]. The decline of ovarian follicle numbers has been studied for decades, and it has been described as potentially involving “pure chance” [4] and a “randomized stochastic event, similar to radioactive decay” [5].

Stochastic modeling has played an important role in understanding follicle loss during ovarian aging, dating back to the seminal 1976 model of Faddy, Jones, and Edwards [6] and the 1983 model of Faddy, Gosden, and Edwards [7] (Edwards later won the Nobel Prize in Physiology or Medicine for developing in vitro fertilization). Such stochastic models have been used to understand the timing of both menopause [8, 9] and the preceding menopausal transition [10] across populations of women. Furthermore, stochastic modeling has recently emerged as an important tool for studying clinical interventions aimed at delaying menopause, including ovarian tissue cryopreservation [11], pharmacological or biologic agents which slow follicle loss [12–14], and therapies which “boost” the number of autologous primordial follicles [14,15]. These clinical applications for women’s health and reproduction underscore the need for scientific rigor in mathematical modeling and analysis of ovarian aging and menopause timing.

In this paper, we analyze a recent model of ovarian aging and menopause timing, which we refer to as the MTK model [16]. In Ref. [16], the authors make the following four claims.

i. The MTK model “predicts the average age of menopause across geographically diverse human populations.”
ii. The MTK model is a “novel theoretical framework.”
iii. The MTK model yields a “universal relation between the initial follicle reserve, the depletion rates, and the threshold that triggers menopause.”
iv. The menopause age distribution “might be a result of a precise regulation due to the synchronization of transitions between different stages of follicles.”

Our investigation of the MTK model is roughly organized around these four claims.

We first show that claim (i) results from choosing parameter values that are orders of magnitude outside of physiological ranges. The error appears to stem from misinterpreting the number of female participants in prior studies as the number of ovarian follicles. Correcting this error invalidates claim (i).

Regarding claim (ii), we use stochastic analysis to put the MTK model in the same framework of previous models [9, 17–19]. Furthermore, we show that the MTK model is a reparameterization of a 1995 model of Faddy and Gosden [17]. Putting the MTK model in this prior framework reveals that menopause age is controlled by the slowest ≈ 0.3% of follicles to exit the ovary, which enables the use of extreme value theory.

Regarding claim (iii), we prove general mathematical results in extreme value theory and use them to obtain an explicit formula that predicts the expected menopause age in the MTK model. This new formula is orders of magnitude more accurate than the heuristic formula posited in Ref. [16] (the formula in Ref. [16] has errors of up to several years). In addition to greater accuracy, the extreme value theory approach is applicable to a broad class of menopause models, which contrasts the “universal” formula posited in Ref. [16]. We further apply our extreme value theory results to the MTK model to obtain explicit estimates of the full probability distribution and all the moments of the menopause age for any given physiological parameters, and we demonstrate their accuracy with stochastic simulations.

Finally, while claim (iv) is an interesting hypothesis, we use extreme value theory to argue that the “narrow distribution” of menopause ages mentioned in Ref. [16] is a generic feature of combining physiological parameter ranges with the modeling framework used in Refs. [9,16–19]. Furthermore, we show how this “narrow distribution” results from solely considering variability from stochastic follicle behavior, and that incorporating the empirical population distribution of initial follicle supply into the MTK model yields a menopause age distribution reminiscent of the distribution observed across populations of women.

## 2 Methods

### 2.1 MTK model

We now describe the model in Ref. [16], which we refer to as the MTK model. The MTK model [16] tracks a population of *N* follicles that irreversibly advance through a chain of *m* ≥ 1 discrete states before ultimately exiting the ovarian reserve upon egress from discrete state *m*. The model is analyzed in terms of the following system of *m* linear ordinary differential equations (ODEs) (see equation (2) in [16]),

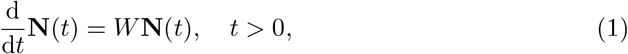

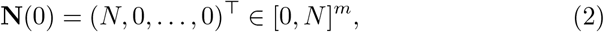

where the vector

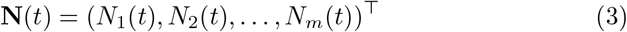

tracks the expected number of follicles in each of the *m* states, and the matrix 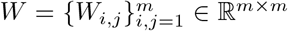 contains the transition rates between states (*W*_*i*,*j*_ is the transition rate from state *j* to *i*). Follicles transition irreversibly from state *i* to state *i* + 1 at rate *λ*_*i*_ *>* 0 for *i* = 1, … , *m* (where state *m* + 1 consists of exited follicles), and thus *W*_*i*,*j*_ = *λ*_*j*_ if *i* = *j* + 1, the diagonal entries of *W* are such that the first *m* − 1 columns of *W* have zero column sums, *W*_*m*,*m*_ = − *λ*_*m*_, and all other entries are zero. The initial condition (2) enforces that all *N* follicles start in state 1.

The discrete state of the continuous-time Markov chain underlying the ODEs in (1) can be represented by the (*m* + 1)-dimensional vector,

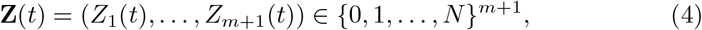

where *Z*_*i*_(*t*) ∈ {0, 1, … , *N*} is the number of follicles in state *i* at time *t* (i.e. *N*_*i*_(*t*) = 𝔼[*Z*_*i*_(*t*)] for *i* = 1, … , *m*) where state *m* + 1 consists of follicles which have exited the ovarian reserve. For each *i* ∈ {1, … , *m*}, each follicle in state *i* moves to state *i* + 1 at rate *λ*_*i*_. Hence, the vector **Z**(*t*) jumps according to

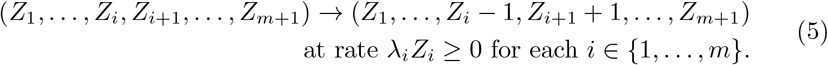

The vector **Z**(*t*) evolves on the state space of (*m* + 1)-tuples of natural numbers whose components sum to *N* , which we denote by 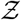,

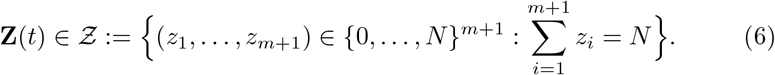

Following [9,17], Ref. [16] defined the menopause age as the first time *T* that only *k* follicles remain in the ovary,

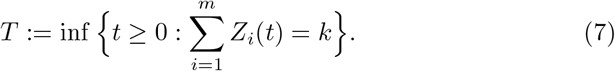

Ref. [16] used the Gillespie algorithm [20] to simulate paths of **Z**(*t*) in (4)-(5) to generate samples of the stochastic time *T* in (7). Ref. [16] assumed that all transition rates are identical,

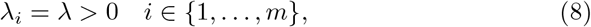

and focused on the case of *m* = 2 states.

### 2.2 Probabilistic analysis

We now present an alternative, probabilistic analysis of the MTK model. Consider a single follicle and let *X*(*t*) ∈ {1, 2, … , *m, m* + 1} denote its discrete state at time *t* ≥ 0, where state *m* + 1 describes a follicle which has exited the ovarian reserve. Suppose the continuous-time Markov chain *X* starts in state *X*(0) = 1 and jumps from state *i* to state *i* + 1 at rate *λ*_*i*_ for *i* = 1, … , *m*. The probability distribution of *X*(*t*) is the vector

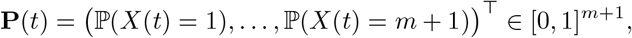

which satisfies the Kolmogorov forward equation [21] (sometimes called the chemical master equation [22, 23]), which is the following system of *m* + 1 linear ODEs,

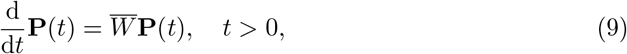

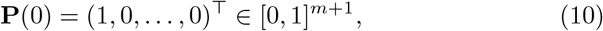

where 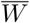 is the (*m* +1) ×(*m* +1)-dimensional matrix with entries 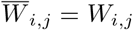 if *i* ≤ *m* and 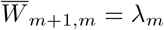, and all other entries of 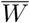 are zero. That is, *X* is a continuous-time Markov chain with infinitesimal generator matrix given by the transpose of 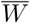 [21].

If 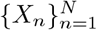 are *N* independent and identically distributed (iid) realizations of *X*, then it follows immediately that **Z** in (4)-(5) can be constructed by defining its *i*-th component, *Z*_*i*_(*t*), as the number of realizations of *X* such that *X*(*t*) = *i*. That is, if |*A*| denotes the cardinality of a set *A*, then

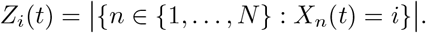

The upshot of this construction is that rather than studying the process **Z**(*t*) in (4)-(5) (which evolves on the very large state space 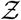 in (6)), we can instead study the process *X*(*t*) (which evolves on the simple state space {1, … , *m* + 1 }).

In addition, the irreversible transitions of *X* further simplify the analysis. In particular, if *τ* denotes the exit time from the ovary of a single follicle,

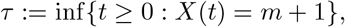

then it follows from (9) and Markov chain theory [21] that *τ* is the sum of *m* independent exponential random variables with rates *λ*_1_, … , *λ*_*m*_, i.e.

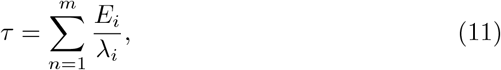

where 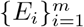 are iid unit rate exponential random variables. For the case of identical transition rates in (8), the exit time *τ* is thus an Erlang random variable with shape *m* and rate *λ*, which means

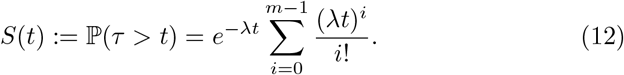

Therefore, the menopause age *T* in (7) is determined by the (*k* +1)-st slowest follicle to exit the ovary, and can thus be written as

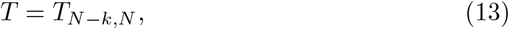

where *T*_1,*N*_ ≤ *T*_2,*N*_ ≤ … ≤ *T*_*N*,*N*_ are the order statistics of *N* iid realizations 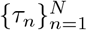 of *τ* in (11),

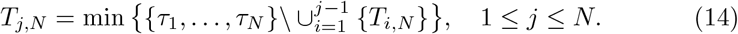

That is, *T* is the (*N* − *k*)-th order statistic of *N* iid Erlang random variables.

### 2.3 General extreme value theory

If *S*(*t*) = ℙ(*τ > t*) denotes the distribution of the stochastic time that a single follicle exits the ovarian reserve, then the independence of *τ*_1_, … , *τ*_*N*_ implies that the distribution of the menopause age *T* = *T*_*N−k*,*N*_ is given exactly by

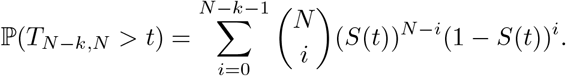

This expression is unwieldy for the parameter regime of interest in which 1 ≪ *k* ≪ *N* . However, this parameter regime means that it is the extreme outlier exit times which determine menopause age, which enables extreme value theory. The basic idea is that the parameter regime 1 ≪ *k* ≪ *N* implies that menopause is triggered by the extreme outlier follicles that are the slowest to exit the ovary. For instance, typical parameters are *k* = 10^3^ and *N* = 3 × 10^5^, in which case menopause is triggered by the slowest 0.3% follicles to exit the ovary.

We now state and prove some general mathematical results in extreme value theory. Let {*τ*_*n*_}_*n≥*1_ be an iid sequence of random variables with common survival probability (i.e. complementary cumulative distribution function), *S*(*t*) = ℙ(*τ > t*). For each *N* ≥ 1 and *j* ∈ {1, … , *N* }, let *T*_*j*,*N*_ denote the *j*-th smallest value in the finite sequence 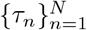. That is, *T*_*j*,*N*_ is the so-called *j*-th order statistic and is defined precisely in (14).

Notice that *T*_*N−k*,*N*_ is the (*k* + 1)-st largest value in 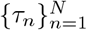 . We are interested in estimating the probability distribution and moments of *T*_*N−k*,*N*_ for large *N* . This distribution depends on the large *t* behavior of *S*(*t*). We are interested in the case that *S*(*t*) decays exponentially, possibly with a power law prefactor,

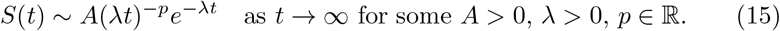

If we can calculate the inverse of *S*(*t*) (denoted *S*^*−*1^), then the following definition is useful for describing *T*_*N−k*,*N*_ for large *N* ,

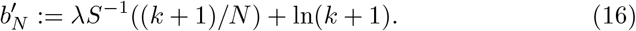

Alternative choices of 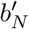 are given in Remark 1 below.

#### Theorem 1.

*Assume S*(*t*) *is a continuous function satisfying* (15). *Assume that* {*b*_*N*_ }_*N≥*1_ *satisfies*

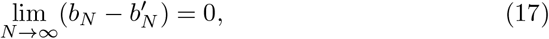

*where* {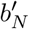}_*N ≥*1_ *is defined in* (16).

*Then for any fixed integer k* ≥ 0, *we have the following convergence in distribution*,

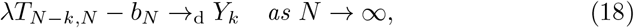

*where Y*_*k*_ *has distribution*

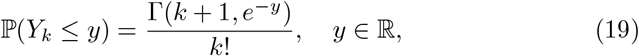

*where* 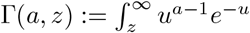 d*u denotes the upper incomplete gamma function.*

#### Remark 1.

Theorem 1 also holds with 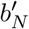 replaced by

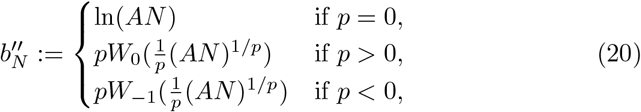

where *W*_0_(*z*) denotes the principal branch of the Lambert W function and *W*_*−*1_(*z*) denotes the lower branch [24]. Theorem 1 also holds with 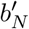 replaced by

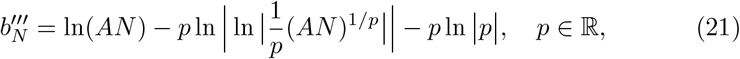

where 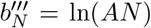 if *p* = 0. See Theorem 1 in [18] for a proof of these two results.

While Theorem 1 describes the probability distribution of *T*_*N−k*,*N*_ for large *N* , the following result gives the moments of *T*_*N−k*,*N*_ for large *N* .

#### Theorem 2.

*If the assumptions of Theorem 1 hold, then for each moment m* ∈ {1, 2, … } *and any fixed k* ≥ 0, *we have that*

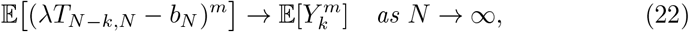

*where Y*_*k*_ *has the distribution in* (19).

#### Remark 2.

Under the assumptions of Theorem 1, it follows immediately from (22) that the mean of *T*_*N−k*,*N*_ has the following asymptotic expansion,

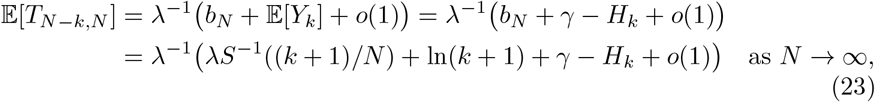

where *γ* ≈ 0.5772 is the Euler-Mascheroni constant, and 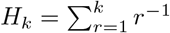 is the *k*-th harmonic number. Furthermore, combining the following expansions,

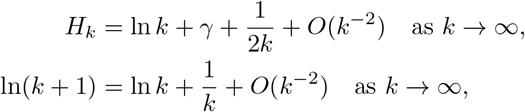

Theorem 2 implies that we can approximate the mean of 𝔼[*T*_*N−k*,*N*_ ] by

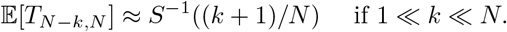

In addition, the variance of *T*_*N−k*,*N*_ has the following asymptotic expansion,

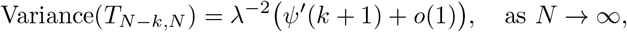

where 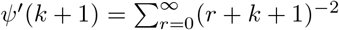 is the first order polygamma function.

### 2.4 Applying extreme value theory to the MTK model

For the MTK model [16] described in sections 2.1-2.2 above, the survival probability of *τ* is given in (12), which for *m* = 2 states reduces to

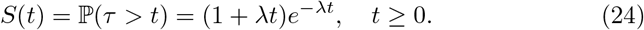

Hence, *S*(*t*) in (24) satisfies (15) with *p* = −1 and *A* = 1. Calculating 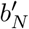 in (16) requires inverting *S*(*t*) in (24), i.e. solving the following equation for *t >* 0,

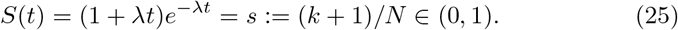

Setting

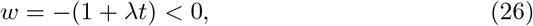

we obtain

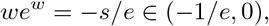

and therefore *w* = *W* (−*s/e*) *<* 0, where *W* denotes either the upper branch, *W*_0_, or the lower branch, *W*_*−*1_, of the Lambert W function [24]. Solving (26) yields 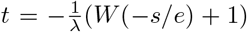. Since we require that *t >* 0, we take the upper branch to obtain the following unique positive solution to (25),

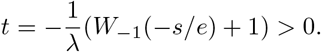

Hence, 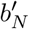 in (16) for the MTK model is

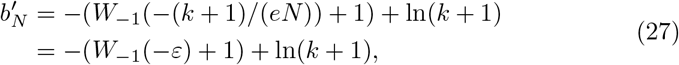

where we have set

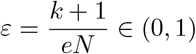

to simplify notation.

Combining Remark 2 with (27) yields the following extreme value theory approximation of the mean menopause age in the MTK model,

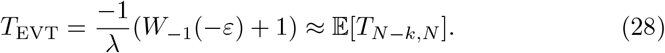

Note that *T*_EVT_ satisfies the following equation,

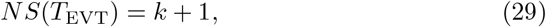

where *S*(*t*) = (1 + *λt*)*e*^*−λt*^. The relation in (29) is intuitive since *NS*(*t*) is the expected number of follicles in the ovarian reserve at age *t* ≥ 0.

We note that retaining the *λ*^*−*1^(ln(*k* + 1) + *γ* − *H*_*k*_) = *O*(1*/*(*λk*)) term in (23) yields an even more accurate approximation, which we denote by *T*_EVT,2_,

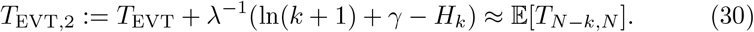

## 3 Results

### 3.1 Correcting the analysis of geographically diverse human populations

A central result of Ref. [16] is based on an error. In Ref. [16], this result is referred to as “Our model accurately predicts the average age of menopause across geographically diverse human populations.” The relevant section in Ref. [16] is entitled “Predictive analysis of menopause timing across geographical regions” and the results are shown in Fig. 6 of Ref. [16].

The result relies on erroneous values of a woman’s follicle supply at birth, which we denote by *N* (denoted *N*_0_ in [16]). Physiological values of *N* are roughly 10^5^ to 10^6^ [25–29]. Table 1 shows some values of *N* used in [16], which are orders of magnitude too small. The error is that these values were taken from what is denoted by “*N* “ in Fig. 4 in Ref. [30], which were in turn taken from Fig. 2 in Ref. [31]. However, these values are not initial follicle supply, but rather the number of women in the study.

**Table 1:** Erroneous values of initial follicle supply used in Ref. [16]. Numbers shown as initial follicle supply are actually the total sample of women providing data on their age at final menses taken from Refs. [30, 31].

| Geographical region | Initial follicle supply |
| --- | --- |
| Africa | 1,175 |
| Asia | 39,158 |
| Australia | 9,268 |
| Europe | 18,962 |
| Latin America | 18,073 |
| Middle East | 7,733 |
| USA | 15,690 |

For example, the value *N* = 1175 for Africa stems from Fig. 2 in Ref. [31] in which 123 women participated in a study in Ghana, 563 women participated in a study in Nigeria, and 489 women participated in another study in Nigeria, and

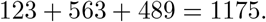

As detailed in section A.1 of the Appendix, the analogous error was made for the six other geographical regions considered in Ref. [16]. In addition, choices of follicle maturation rate *λ* on the order of 10^*−*2^*/*year were used in Ref. [16], which are about an order of magnitude below the physiological range [19].

To summarize, the predictions of Ref. [16] regarding menopause across human populations are based on misinterpreting data and are therefore unfounded.

### 3.2 Unifying prior models

#### 3.2.1 iid follicle exit times

Ovarian aging has been understood in terms of the loss of a dormant ovarian reserve of primordial follicles (PFs). Ref. [16] follows several prior works [6, 7, 9–11, 17–19] (dating back to 1976 [6]) by employing a stochastic model of PF loss in which each of the *N* ≫ 1 PFs present at birth exits the ovarian reserve at independent and identically distributed (iid) random times. Specifically, if *τ*_1_, … , *τ*_*N*_ denote the times that each PF exits the reserve, then these works assume that [6, 7, 9–11, 16–19]

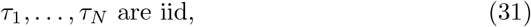

with common probability distribution

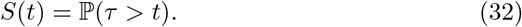

#### 3.2.2 Menopause time

Ref. [16] follows Refs. [9, 18, 19] by connecting the PF decay model in (31)-(32) to menopause timing by assuming that menopause occurs when the number of PFs in the reserve reaches a threshold of *k* PFs, where

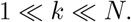

Mathematically, the menopause time is thus the stochastic time *T* in which only *k* PFs remain in the reserve. Precisely, the menopause time *T* is the (*k* + 1)-st slowest follicle exit time [18],

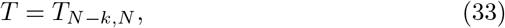

where *T*_1,*N*_ ≤ *T*_2,*N*_ ≤… ≤ *T*_*N*,*N*_ are the order statistics of *τ*_1_, … , *τ*_*N*_ in (31)-(32) (defined precisely in (14)).

#### 3.2.3 Choosing functional forms and parameters

The modeling framework in (31)-(33) requires choosing *N* , *k*, and *S*(*t*) = ℙ(*τ > t*). The menopause threshold *k* can be chosen by comparing follicle data and menopause age data. Since the expected number of follicles remaining in the reserve at age *t* ≥ 0 is *NS*(*t*), both *N* and *S*(*t*) can be parameterized by fitting follicle data. However, before parameterizing *S*(*t*), one must choose its functional form.

Ref. [16] follows Refs. [6, 7, 9, 17] by assuming that *S*(*t*) = ℙ(*τ > t*) is the distribution of a first passage time *τ* of a Markov process. In Refs. [6, 7, 16, 17], the Markov process represents discrete stages of follicle development.

Faddy and Gosden [17] assumed that each follicle stochastically advances through a chain of *m* = 3 discrete states, as depicted in Figure 1. These states correspond to the following biophysically distinct follicle classes (according to the follicle classification system of type B, B/C+C, and D+ [1, 32]). State 1 are unilaminar follicles with a single layer of squamous pregranulosa cells. State 2 are unilaminar follicles consisting of one layer of cuboidal granulosa cells, with or without some squamous cells in the same layer. State 3 are growing follicles with an enlarged oocyte and two or more layers of granulosa cells. The follicle starts in state 1, jumps from state *i* to *i* + 1 at rate *λ*_*i*_, and can exit the reserve at rate *µ*_*i*_ ≥ 0 from state *i*; see Figure 1. Then, *S*(*t*) = ℙ(*τ > t*) is the distribution of the first time *τ* that the follicle exits the reserve. The parameters *λ*_1_, *λ*_2_, *µ*_1_, *µ*_2_, and *µ*_3_ were chosen in [17] by fitting follicle count data for the aforementioned three follicle classes.

**Figure 1:**
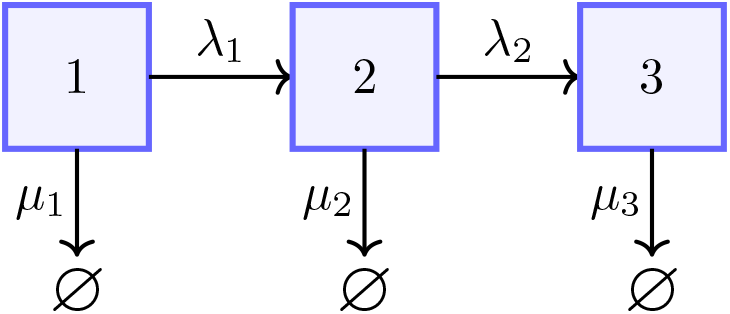
Diagram of an ovarian follicle dynamics model by Faddy and Gosden [17]. Follicles stochastically progress from state *i* to state *i* + 1 at rate *λ*_*i*_ and exit the ovarian reserve from state *i* at rate *µ*_*i*_. This Faddy and Gosden model reduces to the MTK model with *m* = 2 follicle states studied in Ref. [16] upon setting *λ*_1_ = *µ*_2_ = *λ >* 0 and *µ*_1_ = *λ*_2_ = *µ*_3_ = 0.

Setting

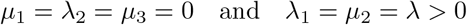

in the Faddy and Gosden model [17] yields the MTK model with *m* = 2 studied in Ref. [16]. The *m* = 3 states in Ref. [17] and the corresponding rates were chosen from data on three biophysically distinct follicle states. In contrast, the choice of *m* = 2 states in Ref. [16] and the single rate *λ* were chosen as a simple model to fit data on the total number of follicles in the ovarian reserve (i.e. without distinguishing between follicle types).

#### 3.2.4 Fit to follicle count data

The fit of the MTK model to the expected primordial follicle (PF) count, *NS*(*t*), is shown in Figure 2 (red dashed curve). Following Ref. [9], the PF “starting supply” was taken to be *N* = 3.23 × 10^5^, which is the median of the 30 PF counts in Ref. [29] taken from women who were at least 6 months gestation and at most one month post birth. For comparison, the blue solid and green dotted curves show the fit of the models posited in Ref. [9] and Ref. [29], respectively. Measuring fit by the sum of squared differences between the logarithms of a model’s expected PF count and the PF count data, the model in Ref. [29] has the best fit, reflecting that five parameters were chosen in Ref. [29] to fit PF data (the model in Ref. [29] also fit PF data before birth). The fit of the model in Ref. [9] is 3% worse than the model in Ref. [29], reflecting that only three parameters were informed by PF count data. The fit of the MTK model is 9% worse than the model in Ref. [29], reflecting that only two parameters (*N* and *λ*) were informed by PF count data.

**Figure 2:**
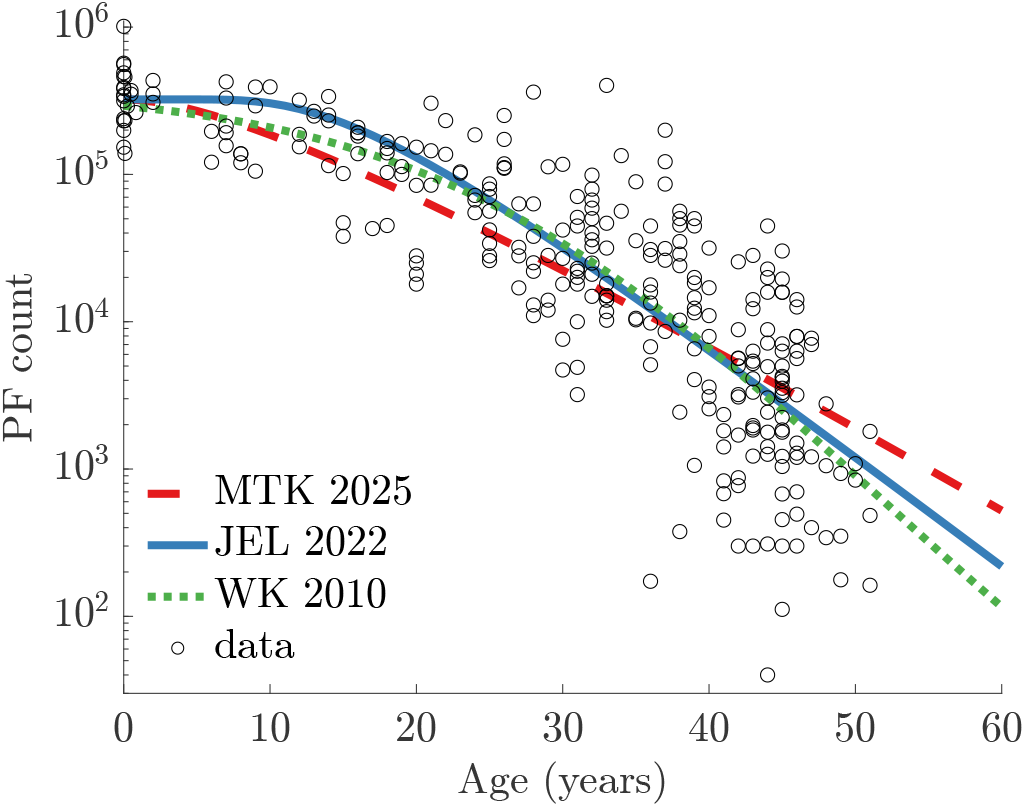
Comparison of PF decay models from Ref. [29] (WK 2010, green dotted), Ref. [9] (JEL 2022, blue solid), and Ref. [16] (MTK 2025, red dashed) to the PF data reported in Ref. [29] (black circles) for 325 women. WK has the best fit to the data, the JEL fit is 3% worse, and the MTK fit is 9% worse, which reflects that WK used 5 parameters, JEL used 3 parameters, and MTK used 2 parameters (*λ* and *N* ). See section 3.2.4 for details.

### 3.3 Improving expected menopause age estimate

As described above, the menopause age in the MTK model is the stochastic time *T* in (33). The following estimate of expected menopause age 𝔼[*T* ] was derived in Ref. [16] using a heuristic argument,

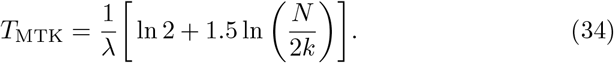

The formula for *T*_MTK_ in (34) is described in Ref. [16] as a “universal relation” between the initial follicle reserve *N* , the depletion rate *λ*, and the threshold *k* that triggers menopause. Similar predictions of menopause age as a function of *N* , *k*, and follicle decay dynamics were previously offered in Refs. [11, 18] for other ovarian aging models.

Using extreme value theory, we derive in Methods the following estimate of expected menopause age 𝔼[*T* ] for the MTK model,

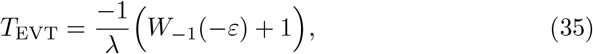

where *ε >* 0 compares the menopause threshold *k* to the initial follicle reserve *N*,

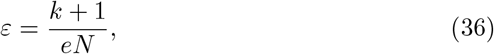

and *W*_*−*1_ denotes the lower branch of the Lambert W function [24]. The Lambert W function in (35) is a standard function that is included in most computational software (it is sometimes called the product logarithm or the omega function). Nevertheless, the Lambert W function can be avoided since the formula in (35) can be approximated to high accuracy in terms of elementary functions. For instance, taking only the first few terms in an asymptotic expansion of *W*_*−*1_ [24] yields the following approximation,

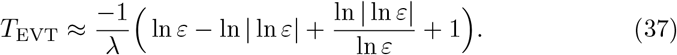

Formulas (35) and (37) agree to within 0.08% for physiologically relevant values of *ε* in (36) (see section A.2 in the Appendix).

Figure 3 compares the *T*_MTK_ formula in (34) and our *T*_EVT_ formula in (35) to stochastic simulations of the MTK model. In this figure, we use parameter values posited in Ref. [16] (namely, *λ* = 0.145*/*year and *k* = 1600). While *T*_MTK_ accurately predicts menopause age 𝔼[*T* ] for *N* ≈ 2 × 10^5^, the formula *T*_MTK_ differs from 𝔼[*T* ] by up to 4 years across the physiological range of *N* = 10^5^ to *N* = 10^6^. In contrast, *T*_EVT_ remains accurate across this range. Indeed, Figure 3B shows that *T*_EVT_ is around 100 to 1,000 times more accurate than *T*_MTK_ as measured by relative error. The relative error in Figure 3B is

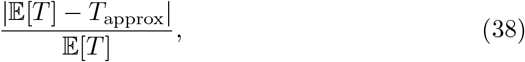

where 𝔼[*T* ] is computed from stochastic simulations and *T*_approx_ is either *T*_MTK_ in (34), *T*_EVT_ in (35), or

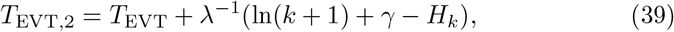

where *γ* ≈ 0.5772 is the Euler-Mascheroni constant and 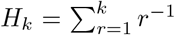 is the *k*-th harmonic number. The formula for *T*_EVT,2_ in (39) is derived in Methods. While Figure 3B shows that *T*_EVT,2_ is even more accurate than *T*_EVT_, the formulas *T*_EVT_ and *T*_EVT,2_ differ by less than a day for physiological parameter values and thus are equivalent for practical purposes.

**Figure 3:**
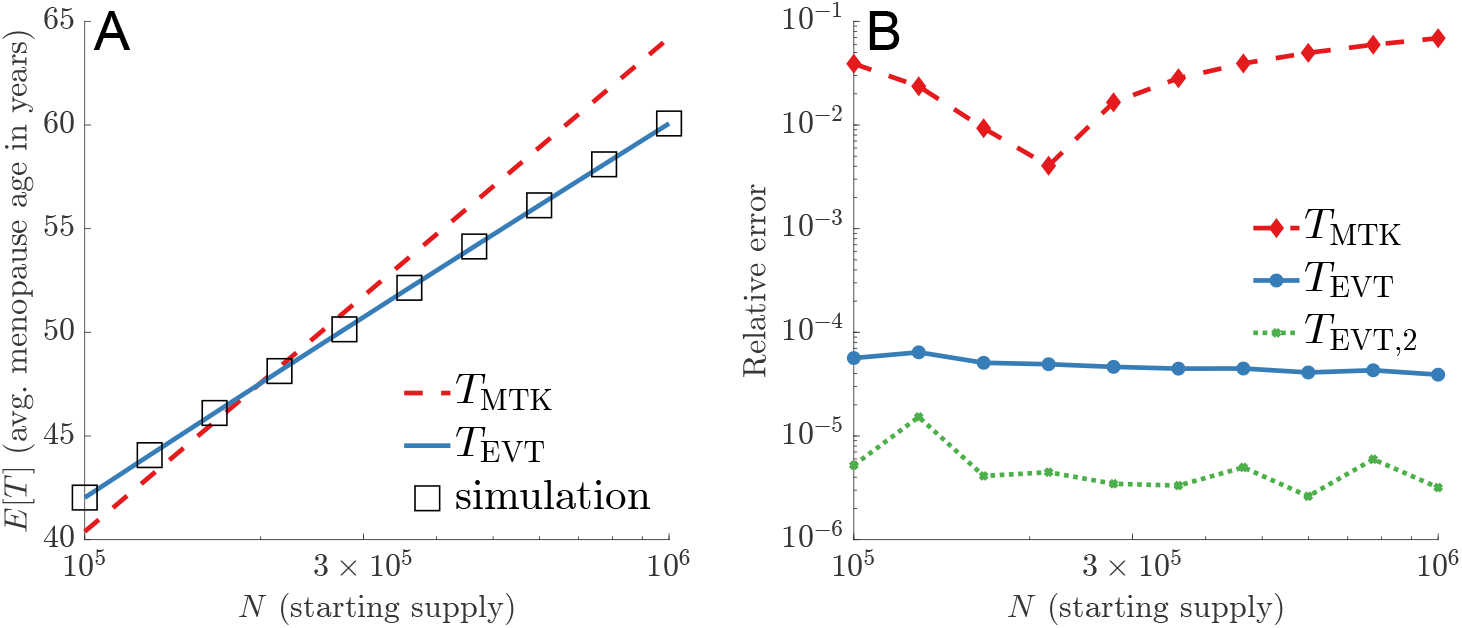
Expected menopause age, 𝔼[*T* ]. Panel A compares *T*_MTK_ in (34) (red dashed) and *T*_EVT_ in (35) (blue solid) to estimates of 𝔼[*T* ] (black squares) computed from 10^6^ statistically exact, independent realizations of *T* in the MTK model. Panel B plots the relative error (38) of the expected menopause age estimates of *T*_MTK_ in (34), *T*_EVT_ in (35), and *T*_EVT,2_ in (39).

In addition to yielding a more accurate estimate of expected menopause age, the extreme value theory approach which yielded *T*_EVT_ in (35) can be considered “universal” in that it can be applied to estimate menopause age in any model which adopts the framework in section 3.2.

### 3.4 Menopause age variability for an individual woman

Section 3.3 above concerns the expected menopause age 𝔼[*T* ] as a function of physiological parameters, and thus does not consider variability. It is important to distinguish between the following two types of variability in the menopause age *T* in the modeling framework of section 3.2:

I. variability for an individual woman, and
II. variability across a population of women.

We consider variability type (I) in this section and type (II) in the following section 3.5 below.

Variability type (I) stems from stochastic follicle exit times and can be understood using extreme value theory. In Theorem 1 in Methods, we rigorously derived the following approximation of the menopause age distribution for an individual woman in the MTK model (i.e. for fixed parameters *N* , *k*, and *λ*),

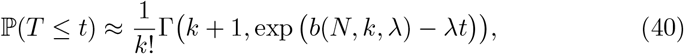

where 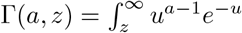 d*u* denotes the upper incomplete gamma function,

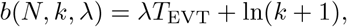

and *T*_EVT_ is in (35).

Figure 4A shows excellent agreement between the analytical approximation in (40) and stochastic simulations of the MTK model. This figure takes *N* = 3.23 × 10^5^ (the median starting supply from the data reported in [29]) and the values *λ* = 0.145*/*year and *k* = 1600 posited in Ref. [16]. Hence, the extreme value theory approach yields not just the mean menopause age in the MTK model, but its full probability distribution. Moreover, the extreme value theory approach is universal in the sense that it can be applied to any model in the modeling framework of section 3.2. Indeed, extreme value theory yielded similar results when applied to the model in Ref. [9] (see figure 1 in section 3 of [18]).

**Figure 4:**
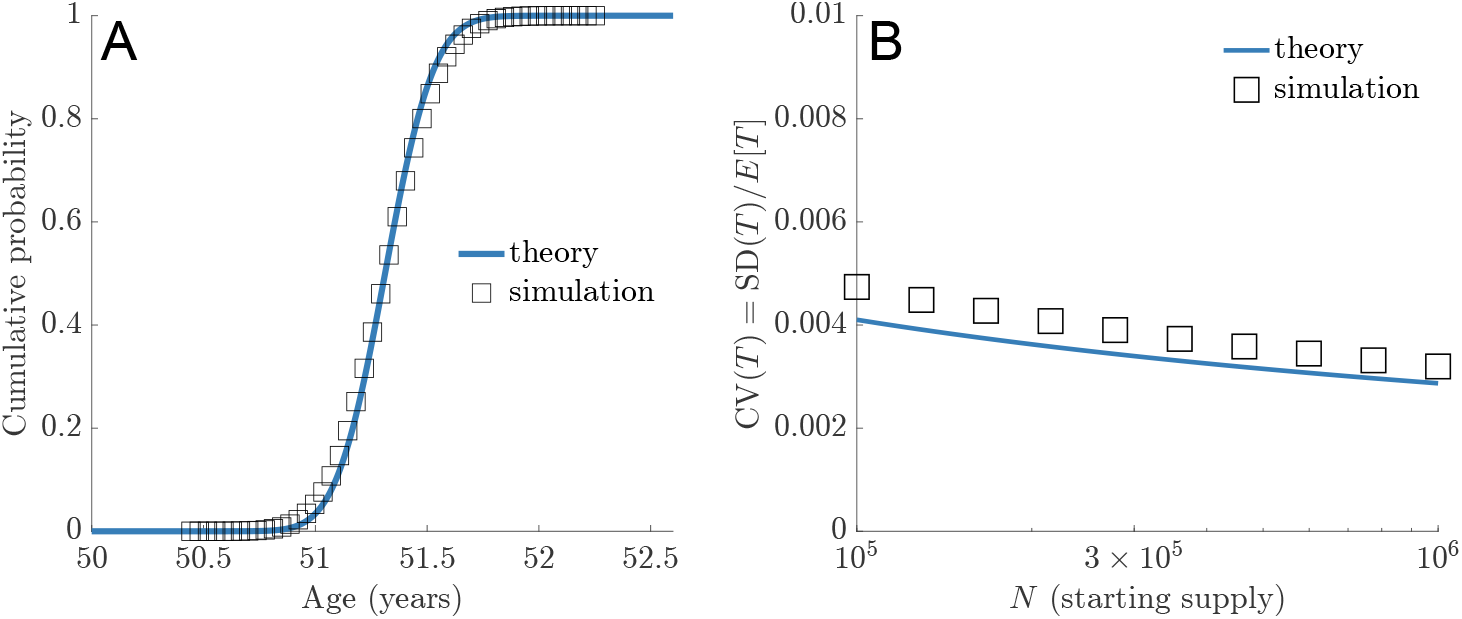
Extreme value theory approximates the full probability distribution of menopause age *T* in the MTK model for fixed parameters. **Panel A:** The blue curve is the formula in (40) and the black squares are computed from 10^6^ statistically exact, independent realizations of *T* in the MTK model. **Panel B**: The blue curve is the formula in (42) and the black squares are computed from 10^6^ statistically exact, independent realizations of *T* in the MTK model. The variability in this figure is “type (I)” rather than “type (II)” (see section 3.4 for details).

Figure 4 shows that type (I) variability is very low. This low variability can be quantified with the following extreme value theory estimate of the standard deviation of the menopause age *T* in the MTK model for fixed *N* , *λ*, and *k*,

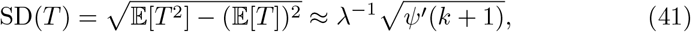

where 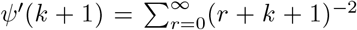 is the first order polygamma function. The approximation in (41) is rigorously justified in Theorem 2 in the Methods. Combining (41) with the extreme value theory estimate of 𝔼[*T* ] in (35) yields the following estimate of the coefficient of variation of *T* ,

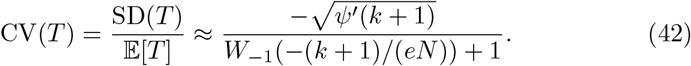

Figure 4B compares the approximation in (42) (solid blue curve) to estimates of CV(*T* ) obtained from stochastic simulations (black squares) with *k* = 1600 and *N* ranges from *N* = 10^5^ to *N* = 10^6^. Figure 4B shows that the formula in (41) agrees with simulations to within around 10% to 13% for these physiologically relevant parameters.

Figure 4B and the extreme value theory estimate in (41) show that the coefficient of variation of *T* is around a mere 0.4%, which translates to a standard deviation of only a few months. This low variability was first noted in Refs. [18, 19] (see section 3 in [18] and figure 5 in [19]). This low variability was later observed in simulations in Ref. [16], and the authors postulated that this “might be a result of a precise regulation due to the synchronization of transitions between different stages of follicles.” We contend that this low variability is instead a generic result of combining the modeling framework of (31)-(33), physiologically relevant parameters, and extreme value theory. Indeed, Theorem 1 in Methods (and also Theorem 2 in [18]) implies that for any model in the framework of section 3.2 for which *S*(*t*) = ℙ(*τ > t*) decays exponentially at large time,

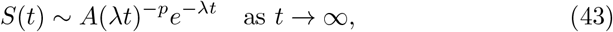

for some *A >* 0, *p* ∈ ℝ, and *λ >* 0, the standard deviation of menopause age, SD(*T* ), must satisfy (41) for large *N* . Physiologically relevant values of the exponential decay rate *λ* in (43) can be obtained by fitting a line to the PF data in Figure 2 in the few decades before age 50 years (note the logarithmic vertical axis). Indeed, prior estimates of the exponential decay rate *λ* in (43) range from *λ* = 0.3*/*year in Ref. [17], to *λ* = 0.24*/*year in Ref. [33], to *λ* = 0.17*/*year in Ref. [19], to *λ* = 0.145*/*year in Ref. [16]. Furthermore, estimates of the menopause threshold range from around *k* = 500 to *k* = 1600 [8, 9, 16, 19, 29, 33]. Plugging these ranges of *λ* and *k* into (41) yields the following range for the standard deviation of the menopause age *T* ,

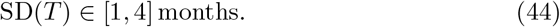

**Figure 5:**
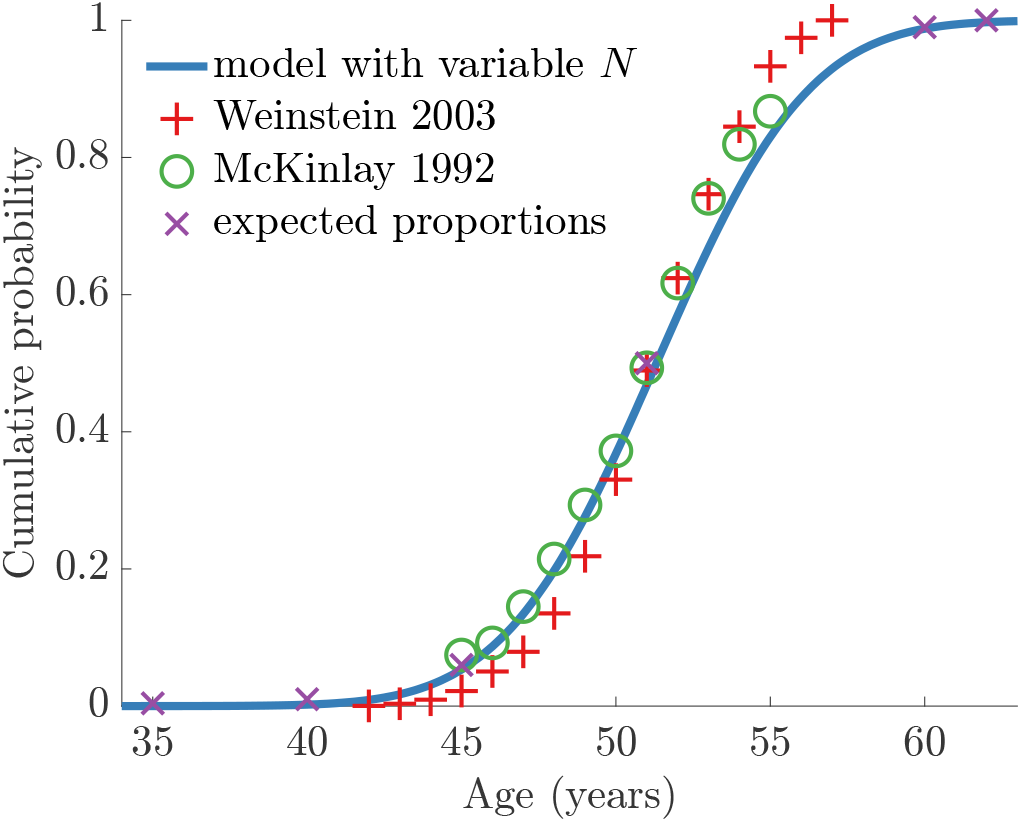
Simulated menopause age distribution across a population of women resembles empirical distributions [10, 37, 38]. See section 3.5 for details.

While this argument relies on the approximation (41), which requires large *N* for accuracy, the agreement between this extreme value theory and simulations in Figure 4 for the MTK model and also figure 5 in Ref. [19] for the model of Ref. [9] suggests the validity of this argument for physiologically relevant parameters.

### 3.5 Menopause age variability across a population

The menopause age standard deviation of a few months in (44) only measures variability stemming from the stochastic behavior of individual follicles. The actual menopause age standard deviation observed across a population of women is at least four years [34, 35]. Indeed, observed menopause ages span roughly twenty years around the population median of age 51 years,

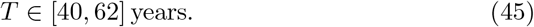

The hundredfold discrepancy between the months in (44) and the decades in (45) can be understood by noting that (44) is for fixed model parameters (i.e. fixed values of *N* , *S*, and *k*). However, it is reasonable to suppose that there is population variability for *N* , *S*, and *k*. For instance, postnatal exposures such as heavy smoking may hasten menopause by around one year [36]. This is an example of an induction of population variability in the decay dynamics *S* and possibly the menopause threshold *k*. Furthermore, a tenfold variation in starting supply *N* across a population is readily visible from the data in Figure 2 at age 0, where *N* can can be seen to vary between roughly *N* = 10^5^ to *N* = 10^6^.

We now show that combining the empirical population distribution of *N* with our analysis of the MTK model yields a menopause age distribution that resembles what has been observed across populations of women. The empirical population distribution of *N* has been estimated by fitting a log normal distribution to the 30 PF counts in Ref. [29] taken from women who were at least 6 months gestation and at most one month post birth [9]. Specifically, this data has been modeled by [9]

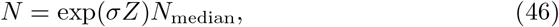

where *N*_median_ = 3.23 × 10^5^, *σ* = 0.5, and *Z* is a standard normal random variable. For (46), the approximate bottom 1% and top 1% of starting supplies *N* are respectively *N* = 10^5^ and *N* = 10^6^.

Figure 5 shows the menopause age distribution across a population of 10^6^ simulated women, where the menopause age *T* for each simulated woman is obtained by first sampling her starting supply *N* from (46) and then calculating *T* via our approximation *T*_EVT_ in (35) (we take *λ* = 0.145*/*year and *k* = 1600 [16]). This calculation relies on the accuracy of our formula for *T*_EVT_ in (35), which is justified since Figure 3 shows that *T*_EVT_ approximates 𝔼[*T* ] to within 0.005%. This calculation also ignores variability in *T* stemming from stochastic follicle exit times, which is justified since we showed in section 3.4 that this accounts for only a few months of variation in *T* .

The simulated menopause age distribution in Figure 5 is similar to menopause age data from Refs. [37, 38] as well as “expected proportions” from the literature [10]. Furthermore, the standard deviation of the simulated menopause age distribution in Figure 5 is 3.9 years, which agrees well with the observed 4 to 4.5 year standard deviation across US women [34]. Similar results have been obtained by combining the starting supply distribution in (46) with other ovarian aging and menopause timing models [9, 10]. While women likely vary in follicle decay dynamics and menopause threshold (*S* and *k* in the modeling framework of section 3.2), this analysis suggests that variations in starting supply *N* may be the dominant source of menopause age variability across a population.

## 4 Discussion

In this paper, we analyzed a recent model of ovarian aging and menopause timing in the context of prior work, dating back to the 1970s and 1980s [6, 7]. We first corrected an error regarding menopause age across human populations. We then used stochastic analysis to show that this MTK model (a) follows Refs. [6, 7, 9–11, 17–19] by assuming that follicles exit the ovary at iid random times (determined by first passage times of Markov processes), and (b) follows Refs. [9,17–19] by assuming that menopause occurs when the number of follicles remaining in the ovary crosses a critical threshold. Putting the MTK model in this prior framework allowed the use of extreme value theory. We proved extreme value theory results and used them to gain detailed insights into the implications of this recent model and also the more general modeling framework of assumptions (a) and (b).

The application of mathematical, and specifically, stochastic models to the biological process of ovarian aging has resulted in mechanistic insights about the process and useful tools for probing the area further. Continued collaborations between mathematicians, biologists, and clinicians promise to improve our understanding of reproductive aging, and to support the development of interventions that can improve fertility outcomes and the ongoing health of women. Further careful and rigorous work by a growing number of interested investigators will help to fulfill this promise.

## Acknowledgments

SDL is supported by the National Science Foundation (NSF DMS-2325258 and NSF CAREER DMS-1944574). JJ is supported by the National Science Foundation (NSF DMS-2325259), CU-Anschutz School of Medicine Funds, and CU-Anschutz Department of Obstetrics and Gynecology Research Funds.

## Data availability statement

Data will be made available on reasonable request.

## A Appendix

### A.1 Erroneous follicle counts

As summarized in the main text, a central claim of Ref. [16] is based on an error. In Ref. [16], this claim is referred to as “Our model accurately predicts the average age of menopause across geographically diverse human populations.” This claim is unfounded since it is based on erroneous data. The purpose of this section is to explain the source of this error in more detail.

The error is that the number of study participants in prior publications was assumed to be the number ovarian follicles. These erroneous values of “*N* “ for seven geographical regions are collected in Table 1 in the main text. The value *N* = 1175 is explained in the main text. The value *N* = 39158 for Asia stems from Fig. 2 in Ref. [31] in which 493 women participated in a study in Japan, 124 women participated in a study in Taiwan, 33054 women participated in a study in China, 2658 women participated in a study in South Korea, 201 women participated in a study in India, 1011 women participated in a study in China, 961 women participated in a study in South Korea, and 656 women participated in a study in Singapore, and

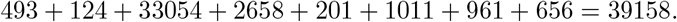

The value *N* = 9268 for Australia stems from Fig. 2 in Ref. [31] in which 7575 women participated in a study in Australia and 1693 women participated in another study in Australia, and

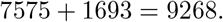

The value *N* = 18962 for Europe stems from Fig. 2 in Ref. [31] in which 2117 women participated in a study in Poland, 490 women participated in a study in Sweden, 3513 women participated in a study in the United Kingdom, 6877 women participated in a study in Czechoslovakia, 1009 women participated in a study in Germany, and 4686 women participated in a study in the Netherlands, and

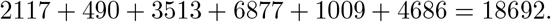

The value *N* = 18073 for Latin America stems from Fig. 2 in Ref. [31] in which 17150 women participated in a study across 15 countries, 472 women participated in a study in Mexico, and 451 women participated in a another study in Mexico, and

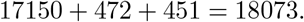

The value *N* = 18073 for the Middle East stems from Fig. 2 in Ref. [31] in which 500 women participated in a study in Iran, 2462 women participated in a study in Tehran, 948 women participated in a study in Iran, 742 women participated in a study in the United Arab Emirates, 1076 women participated in a study in Turkey, 1397 women participated in a study in Iran, 346 women participated in another study in Iran, and 262 women participated in a study in Turkey, and

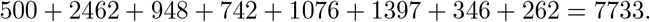

Finally, the value *N* = 15690 for the United States of America stems from Fig. 2 in Ref. [31] in which six studies in the USA are cataloged with respective participant counts of 543, 248, 1825, 1311, 10440, and 1323, and

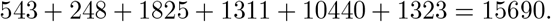

### A.2 Logarithmic expansion of expected menopause age

The Lambert W function *W*_*−*1_ in *T*_EVT_ in (35) is a commonly used mathematical function and is included in most computational software. Nevertheless, *W*_*−*1_ cannot be expressed in terms of elementary functions [24]. If one wants to avoid evaluating the Lambert W function, the following series representation of *W*_*−*1_ can be used [24],

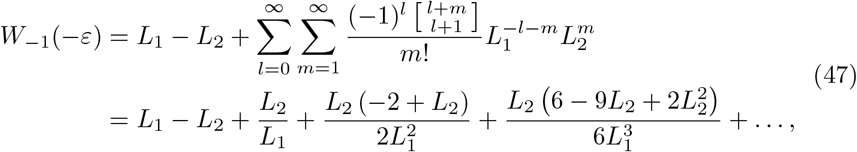

where 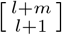 is a non-negative Stirling number of the first kind and

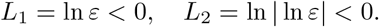

Since *ε* ≪ 1 for physiological parameters, we obtain an accurate approximation to *W*_*−*1_(−*ε*) by taking only the first few terms in (47). In particular, taking the first three terms yields

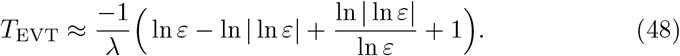

In Figure 6, we plot the relative error in computing *T*_EVT_ in (35) using the Lambert W function compared to using the first one, two, or three terms in the series in (47). Since physiologically relevant values of *N* and *k* are *N* ≥ 10^5^ and *k* ≤ 1600, we typically have

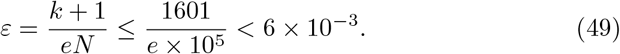

**Figure 6:**
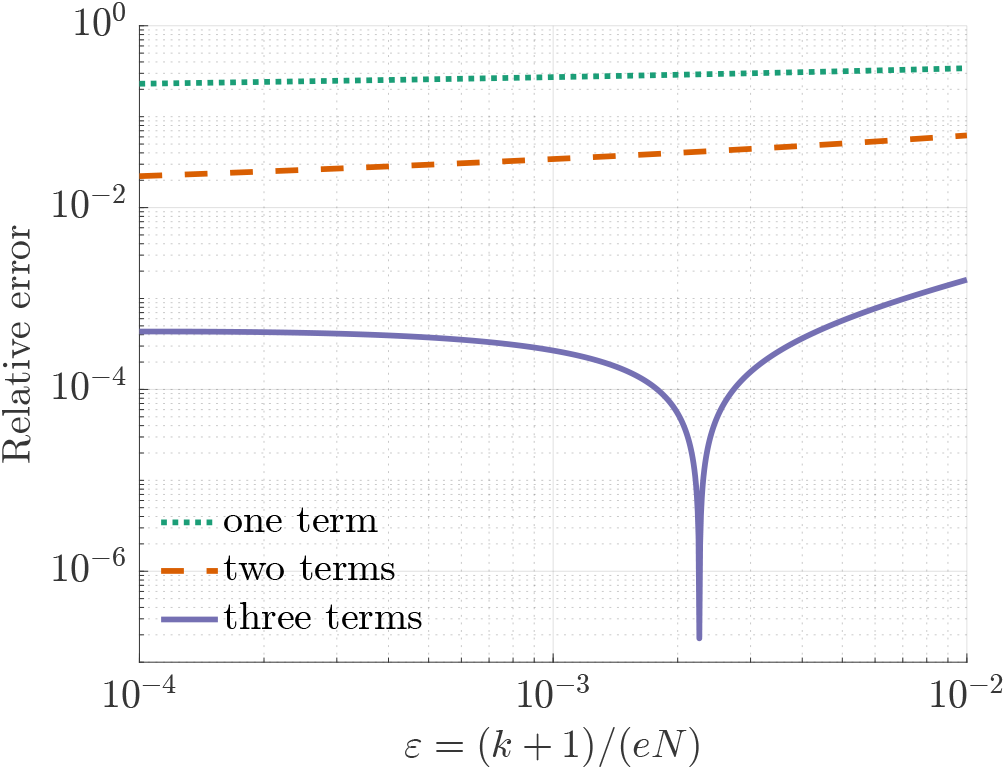
Calculating of *T*_EVT_ in (28) with either the full Lambert W function *W*_*−*1_ or the first one, two, or three terms of the series in (47).

Figure 6 thus shows that the formula in (48) agrees with *T*_EVT_ in (28) to within 0.08% for physiologically relevant values of *ε* (that is, for values of *ε* that satisfy (49)).

### A.3 Proofs of theorems

#### Proof of Theorem 1.

Fix *y* ∈ ℝ. Define

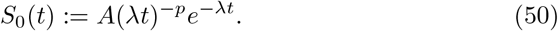

Let *k*^*′*^ = *k* + 1 to simplify notation. The asymptotic equivalence in (15) and the definition of 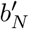 in (16) implies that

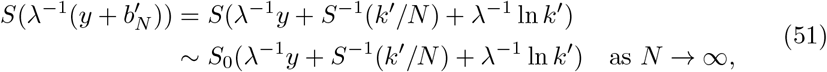

where we have used that (15) implies that

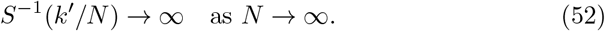

Note that since *S* is a continuous function, *τ* must be a continuous random variable, and therefore *S*(*t*) must be strictly decreasing for sufficiently large *t*, and therefore *S*^*−*1^((*k* +1)*/N* ) exists and 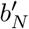 in (16) is well-defined for sufficiently large *N* . The definition of *S*_0_ in (50) implies that

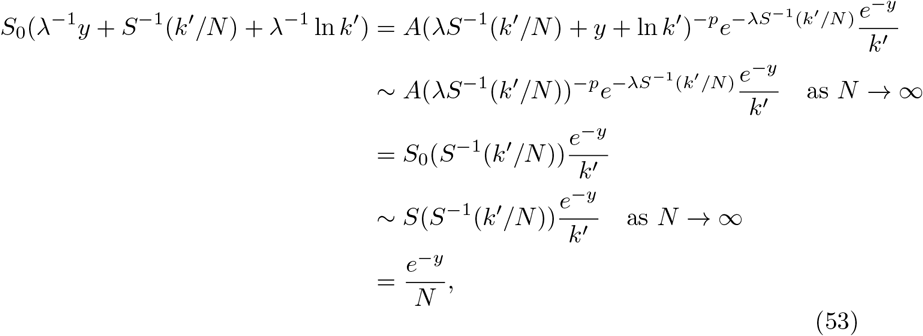

where we have again used (15) and (52). Combining (51) and (53) yields

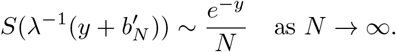

Since lim_*N→∞*_(1 − *N* ^*−*1^*e*^*−y*^)^*N*^ = exp(−*e*^*−y*^), Lemma 1 below yields

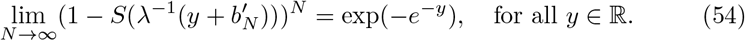

Therefore, since *T*_*N*,*N*_ = max{*τ*_1_, … , *τ*_*N*_ } and {*τ*_*n*_}_*n≥*1_ are iid, we have

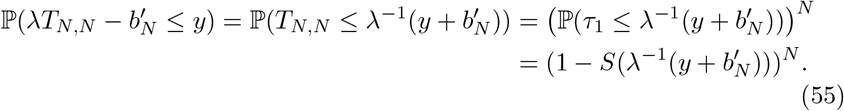

Taking *N* → ∞ in (55) and using (54) implies that (18) holds for *k* = 0 and *b*_*N*_ = 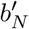 . Theorem 3.4 in [39] then implies that (18) holds for any fixed integer *k* ≥ 1 and *b*_*N*_ = 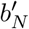. Having proven (18) for *b*_*N*_ = 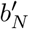, generalizing to any *b*_*N*_ satisfying (17) can be shown using basic properties of convergence in distribution [40].

The following lemma was proven in [18] (see Lemma 9 in section A.1 of [18]).

#### Lemma 1.

*Suppose* {*s*_*N*_}_*N≥*1_ *and* {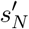}_*N ≥*1_ *are sequences of real numbers satisfying*

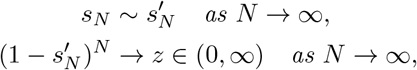

*for some z* ∈ (0, ∞). *Then*

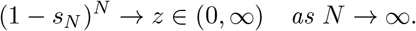

#### Proof of Theorem 2.

Since Theorem 1 holds for any *b*_*N*_ satisfying (17) where 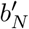 is either defined in (16) or 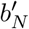 is replaced by 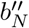 defined in (20), it follows that

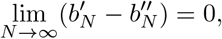

where 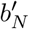 is defined in (16) and 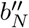 defined in (20). The proof of Theorem 2 above then follows from Theorem 2 in [18].

